# Profiling the effects of rifaximin on the healthy human colonic microbiota using a chemostat model

**DOI:** 10.1101/828269

**Authors:** Ines B. Moura, Anthony M. Buckley, Duncan Ewin, Emma Clark, Suparna Mitra, Mark H. Wilcox, Caroline H. Chilton

## Abstract

Rifaximin is a low solubility antibiotic with activity against a wide range of bacterial pathogens. It accumulates in the intestine and is suitable for prolonged use. Three chemostat models (A, B and C) were used to investigate the effects of three rifaximin formulations (α, β and κ, respectively) on the gut microbiome. Bacterial populations were monitored by bacterial culture and 16S rRNA gene amplicon (16S) sequencing. Limited disruption of bacterial populations was observed for rifaximin α, β and κ. All formulations caused declines in total spores (∼2 log_10_ cfu ml^-1^), *Enterococcus* spp. (∼2 log_10_ cfu ml^-1^ in models A and C, and ∼1 log_10_ cfu ml^-1^ in model B), and *Bacteroides* spp. populations (∼3 log_10_ cfu ml^-1^ in models A and C, and ∼4 log_10_ cfu ml^-1^ in model B). Bacterial populations fully recovered during antibiotic dosing in model C, and before the end of the experiment in models A and B. According to the taxonomic analysis, prior to rifaximin exposure, Bifidobacteriaceae, Ruminococcaceae, Acidaminococcaceae, Lachnospiraceae and Rikenellaceae families represented >92% of the total relative abundance, in all models. Within these families, 15 bacterial genera represented >99% of the overall relative abundance. Overall, the 16S sequencing and culture data showed similar variations in the bacterial populations studied. Among the three formulations, rifaximin κ appeared to have the least disruptive effect on the colonic microbiota, with culture populations showing recovery in a shorter period and the taxonomic analysis revealing the least global variation in relative abundance of prevalent groups.

## Introduction

Rifaximin is an oral antibiotic with low solubility and *in vitro* bactericidal activity reported against a wide range of facultative and obligate aerobes such as, *Escherichia coli, Clostridioides difficile, Staphylococcus* spp., and *Streptococcus* spp., among others [1-3]. Its chemical structure is a derivative of rifamycin and contains a supplementary pyridoimidazole ring that limits the absorption of the drug in the intestine [3]. Rifaximin is considered suitable for prolonged use due to its poor solubility and subsequent low systemic side effects [2, 4]. It is FDA approved for the treatment of traveller’s diarrhoea and irritable bowel syndrome (IBS), and has been shown effective in clinical trials as therapeutic agent in small bowel bacterial overgrowth, inflammatory bowel disease and in *C. difficile* infection [2, 4-8]. Rifaximin is also used to reduce the risk of hepatic encephalopathy in patients with impaired liver function and portosystemic shunting [9].

A decline in colonic bacterial diversity has been reported during rifaximin use, [10-12] with bacterial populations recovering following antibiotic treatment. However, studies have mostly investigated variations in faecal samples from particular patient groups with gastrointestinal or immune diseases [10, 12-14], who may already have a depleted gut microbiota, therefore giving only a partial insight into the effects of rifaximin on the gut microbiome. Sampling during *in vivo* studies is usually done via faecal microbiota profiling which limits the time points available for analysis and does not always reflect disruptive effects in the colon and, providing only partial data on the antibiotic effects in the intestinal microbiome. A continuous *in vitro* model of the human colon has been used before to investigate rifaximin effects in microbiota representative of Crohn’s disease, [15] but sample collection was also limited to pre- and post-antibiotic dosing periods. Understanding the differences in microbiota disruption in healthy and diseased states is of key importance, particularly for antibiotics of prolonged use.

The chemostat gut model used in this study is a clinically reflective representation of the bacterial composition and activities of the human colon, and has been validated against gut contents of sudden death victims [16]. Due to the low solubility of rifaximin, high concentrations can be achieved in the colon, which makes this model ideal to test the disruptive effects of rifaximin on the microbiota [17].The gut model has a proven track record in assessing drug efficacy during pre-clinical and clinical drug development stages. It has been shown to be particularly predictive as an *in vitro* model of *C. difficile* infection (CDI) to evaluate drug propensity to induce CDI, [18-21] with the data correlating well with *in vivo* and clinical trial results [22, 23].

In this study, we used the *in vitro* gut model to investigate the changes in the human healthy intestinal microbiota following instillation of three proprietary rifaximin formulations. Bacterial culture analysis and microbial diversity analysis by 16S rRNA gene amplicon (16S) sequencing were performed to evaluate gut microbiota populations depletion and recovery.

## Materials and Methods

### *In vitro* gut model and experimental design

Three triple-stage chemostat models were assembled as previously described [18, 24], and run in parallel. Briefly, each model consisted of three glass vessels maintained at 37°C and arranged in a weir cascade. Each vessel represents the conditions of the human colon, from proximal to distal, in pH and nutrient availability. The models were continuously fed with complex growth media [18] connected to vessel 1 at a pre-established rate of 0.015h^-1^ and an anaerobic environment was maintained by sparging the system with nitrogen. Sample ports in all vessels allowed sample collection for monitoring of microbiota populations and antibiotic instillation. All models were initiated with a slurry of pooled human faeces from healthy volunteers (n=4) with no history of antibiotic therapy in the previous 6 months, diluted in pre-reduced phosphate-buffered saline (PBS) (10% w/v). Bacterial populations were allowed to equilibrate for 2 weeks (equilibration period) prior to antibiotic exposure (rifaximin dosing period), and to recover for 3 weeks post antibiotic dosing (recovery period), as outlined in Fig. 1. Three proprietary rifaximin formulations, named in this study as alpha-α, beta-β, and kappa-κ (supplied by Teva Pharmaceuticals USA, Parsippany, NJ, USA) were investigated. Model A was inoculated with rifaximin α, model B was inoculated with rifaximin β and model C was inoculated with rifaximin κ. Only one model was used per formulation, as the reproducibility and clinically reflective nature of this system has been shown [16, 21, 23-25]. Due to rifaximin poor solubility, each antibiotic dose was suspended in water and added to vessel 1 of the respective model. Each model was dosed with 400 mg of rifaximin, thrice daily, for 10 days, to replicate the dosing regimen previously used in human clinical trials [4], and the rifaximin concentration that reaches the human colon. No adjustments to the dosing concentration were performed based on the gut model vessel volumes, since the oral administration of a single radiolabelled dose of 400 mg of rifaximin to healthy individuals showed that nearly all rifaximin (∼97%) is excreted in the faeces [2]. Selective and non-selective agars (Table 1) were used for culture profiling of total facultative and obligate anaerobes, Enterobacteriaceae, *Enterococcus* spp., *Bifidobacterium* spp., *Bacteroides* spp., *Lactobacillus* spp., *Clostridium* spp., and total spores, as previously described [18], supported by MALDI-TOF for specific colony/species identification. Bacterial populations in vessel 3 were monitored by inoculating the agar plates in triplicate every other day prior to antibiotic exposure and daily thereafter. The limit of detection for culture assay was established at ∼1.22 log_10_ cfu ml^-1^. Additional samples were taken from vessel 3 of each model to investigate bacterial diversity by 16S sequencing as outlined in Fig. 1. Vessel 3 was particularly investigated due to its microbial richness and representation of the human colon region more physiologically relevant for opportunistic infections [16, 18, 24].

**Table 1.**
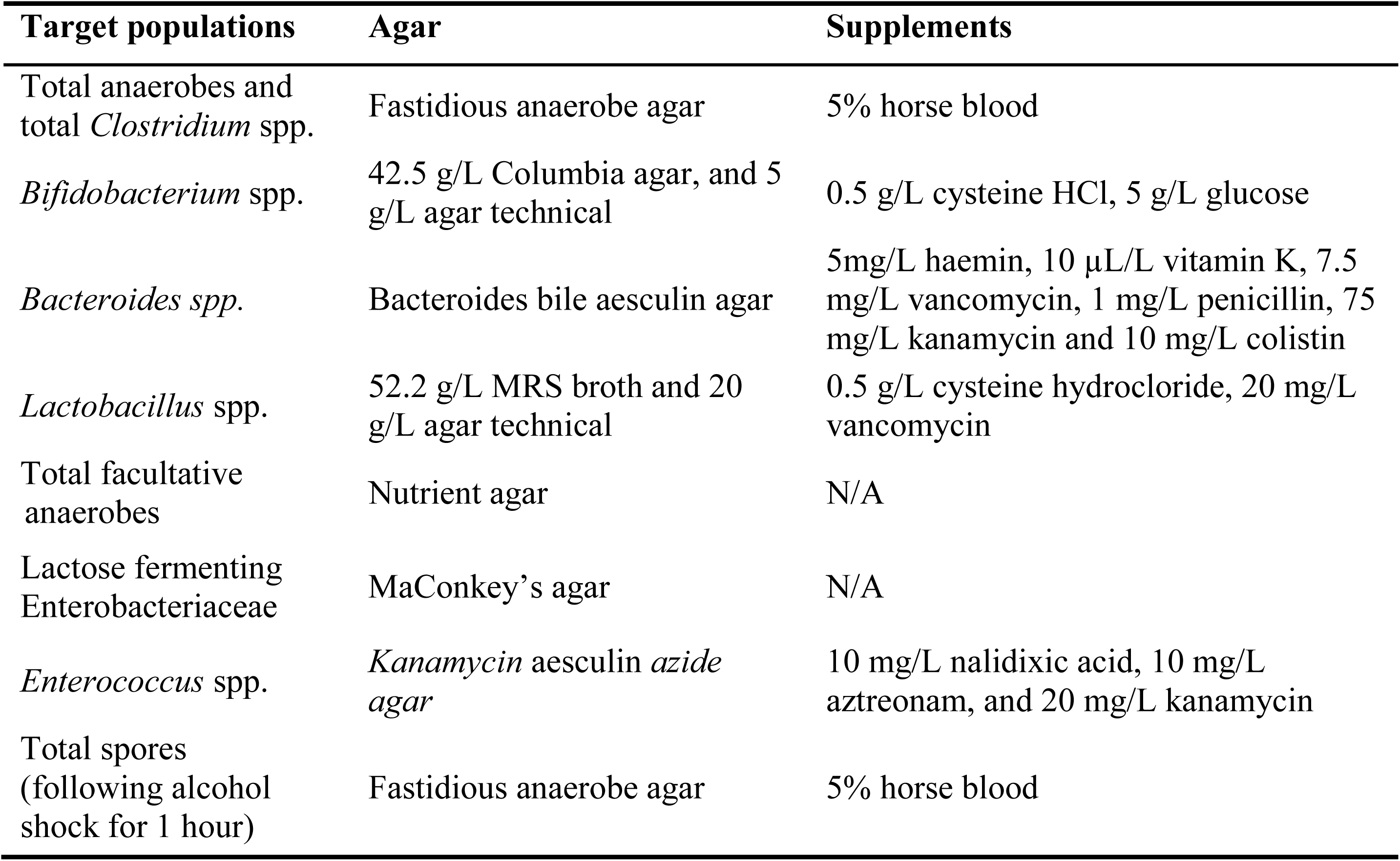
Target populations and agar composition for bacterial ennumeration.

**Figure 1.**
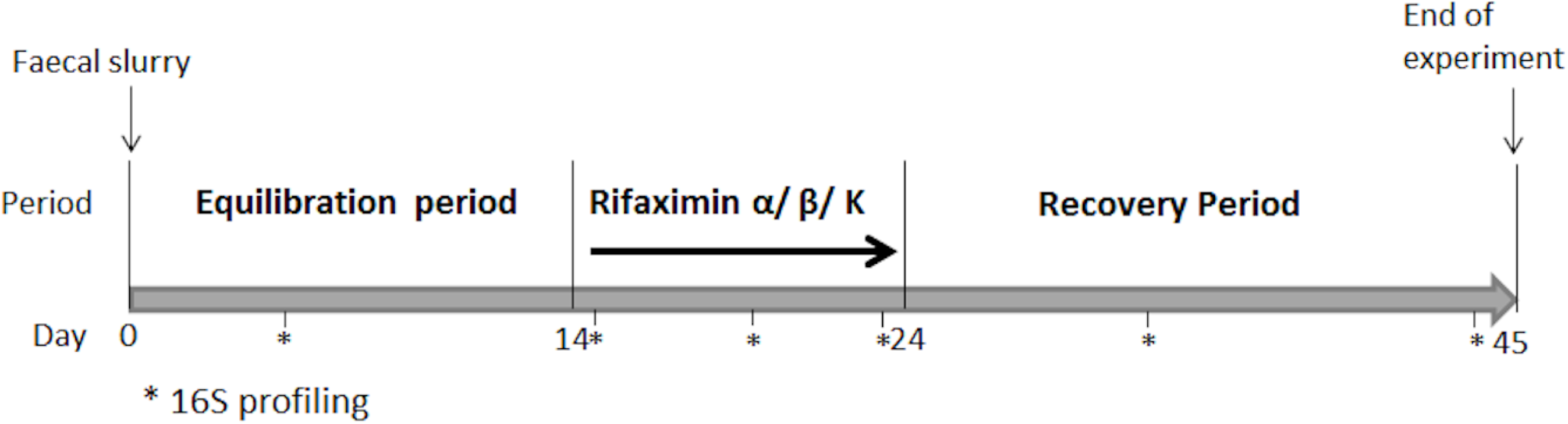
Outline of the gut model experiments with rifaximin α (Model A), rifaxmin β (Model B), or rifaximin κ (Model C). Asterisks indicate the time points (days 5, 15, 20, 23, 33 and 44) of sample collection for DNA extraction and microbial diversity analysis by 16S rRNA gene amplicon sequencing.

### 16S rRNA gene amplification and library preparation

DNA extraction from gut model fluid was performed using the QIAamp DNA Stool Kit with Pathogen Lysis tubes (Qiagen). Samples were pelleted and the supernatant. Protocol was performed according to the manufacturer’s instructions except: a sterile glass bead was added to the lysis tube, homogenisation was performed at 6500 rpm 2×20s using Precellys 24 (Bertin Instruments), and sample clean-up was improved with 20 µg of RNase (Thermo Fisher Scientific). PCR of the variable 4 region (F-5’–AYTGGGYDTAAAGNG–3’, R-5’– TACNVGGGTATCTAATCC–3’), [26] was performed in a 50 µL reaction volume consisting of 40 ng/µL template DNA, 1x Q5 Hot Start High-Fidelity Master Mix (Qiagen), and 25 µM of each primer. Thermal cycler conditions were as follows: denaturing at 98°C for 30s, 30 annealing cycles of 98°C for 5s, 42°C for 10s, 72°C for 20s, and elongation at 72°C for 2 min. Successful amplification was confirmed by gel electrophoresis before samples were cleaned using the MinElute PCR Purification kit (Qiagen). PCR products were quantified and ∼80 ng of dsDNA was carried forward to library preparation using the NEBNext Ultra DNA Library Prep Kit for Illumina and NEBNext Multiplex Oligos for Illumina (New England Biolabs). Following twelve cycles of PCR enrichment, the libraries were cleaned with AMPure Beads (Beckman Coulter), and quality was confirmed by DNA screen tapes (TapeStation, Agilent). Successful libraries were pooled and pair-end sequenced on an Illumina MiSeq platform (2 × 250 bp).

### Bioinformatics analysis and bacterial identification

Sequencing analysis was performed as previously described [27]. Briefly, de-multiplexed FASTQ files were trimmed of adapter sequences using cutadapt [28]. Paired reads were merged using fastq-join [29] under default settings and then converted to FASTA format. Consensus sequences were removed when containing ambiguous base calls, two contiguous bases with a PHRED quality score lower than 33 or a length more than 2 bp different from the expected length of 240 bp. Further analysis was performed using QIIME [30]. Operational taxonomy units (OTUs) were selected using Usearch [31], and aligned with PyNAST using the Greengenes reference database [32]. Taxonomy was assigned using the RDP 2.2 classifier [33]. The files resulting from the above analyses were imported in the Metagenome ANalyzer (MEGAN) for detailed group specific analyses, annotations and plots [34].

### Antimicrobial assay

Bioactive rifaximin concentrations were measured daily during and post antibiotic instillation. Aliquots were collected daily from each vessel of model A, B and C and kept at −80°C until processing. Due to the low solubility of rifaximin, the concentrations of both the solubilised fraction and the concentrations of the antibiotic that remained as suspension in the vessels were measured. Following centrifugation for 10 min at 15,000g, supernatants were removed and 1 ml of methanol was added to each pellet to dissolve any antibiotic powder. An additional centrifugation step was performed to remove cell debris. Concentrations of rifaximin were determined using Wilkins Chalgren agar with *Staphylococcus aureus* as the indicator organism. Assays were performed as previously described [21] in triplicate to assess the concentrations of solubilised antibiotic (supernatants) and the concentrations of antibiotic that remained in suspension (methanol-solubilised powder) in each vessel. The limit of detection for this assay was established at 8 mg l^-1^.

## Results

### Effects of rifaximin α, β and κ on the gut microbiota populations

Three chemostat models (A,B and C) were run in parallel to investigate the effects of three rifaximin formulations (α, β and κ) on the gut microbiota. All models were started with the same slurry and were left without intervention for 14 days to allow bacterial populations to equilibrate. Results for vessel 3 only are shown, as this vessel represents the distal colon, a region of high bacterial diversity and biologically relevant for intestinal diseases associated with disruption of the normal gut microbiota (e.g. CDI) [24]. In all models, microbiota populations were stable prior to antibiotic exposure (Fig. 2 and Fig. S1). Rifaximin α exposure caused a declines in in *Bifidobacterium* spp. and total spores (∼1.5 log_10_ cfu ml^-1^), in *Enterococcus* spp. (∼2 log_10_ cfu ml^-1^), in *Bacteroides* spp. (∼3 log_10_ cfu ml^-1^) and, in *Clostridium* spp. (∼1 log_10_ cfu ml^-1^) populations (Fig. 2a and Fig. S1a). Total spores, *Bifidobacterium* spp. and *Bacteroides* spp. populations recovered during antibiotic instillation period. *Enterococcus* spp. recovered to pre-antibiotic levels by the end of the experiment.. Following rifaximin α instillation, *Clostridium* spp. populations remained at ∼6 log_10_ cfu ml^-1^ throughout the experiment (Fig. S1a).

**Figure 2.**
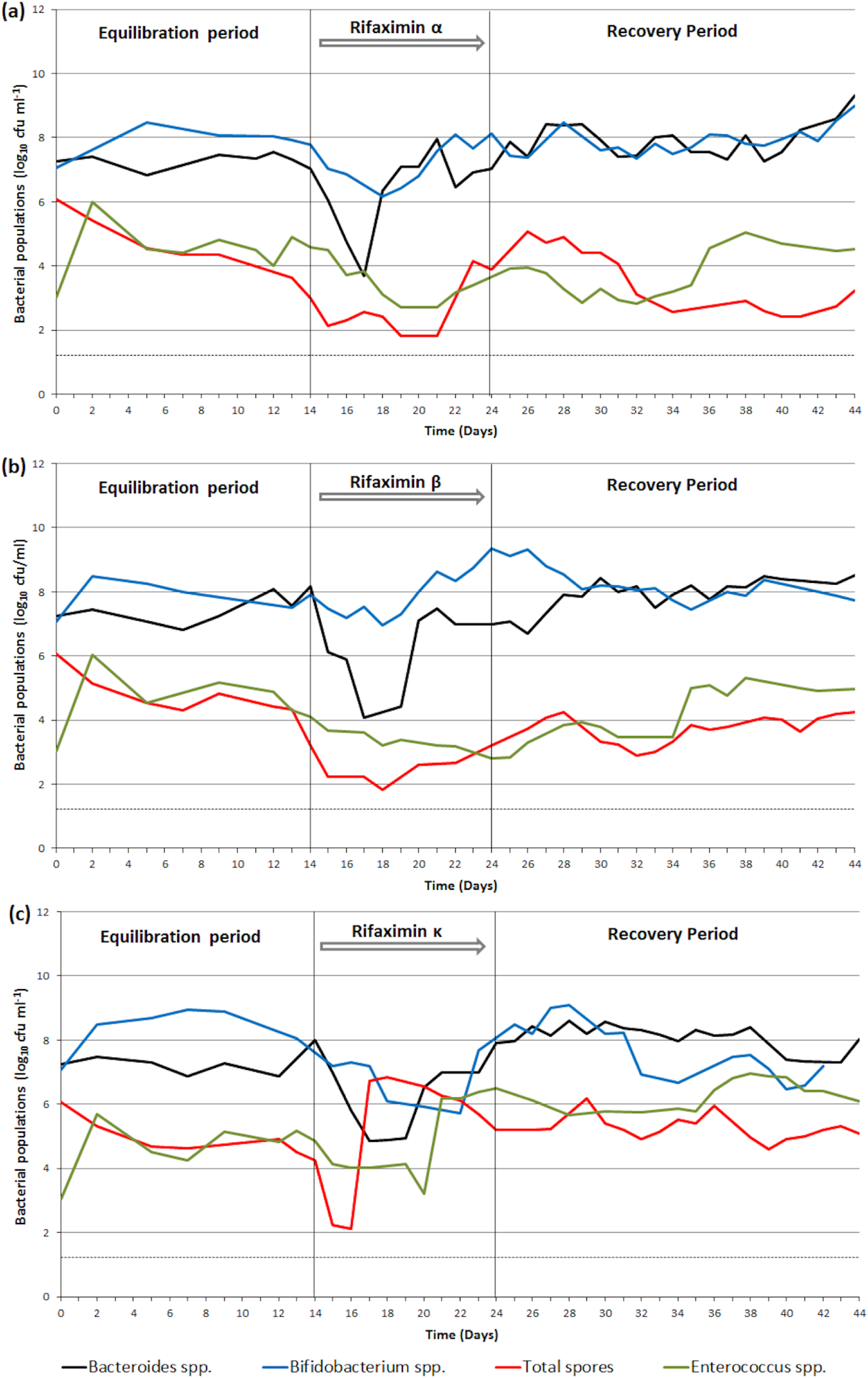
Mean (log_10_ cfu ml^-1^) gut microbiota populations that showed variations during antibiotic instillation in vessel 3 of (a) model A (rifaximin α dosing), (b) model B (rifaximin β dosing), (c) model C (rifaximin κ dosing). Horizontal dotted line marks the limit of detection for culture assay (∼1.2 log_10_ cfu ml^-1^).

In model B, effects of exposure to rifaximin β were similar to those of rifaximin α (Fig. 2b and Fig. S1b). Declines were observed in *Enterococcus* spp. (∼1 log_10_ cfu ml^-1^), total spores (∼2 log_10_ cfu ml^-1^), *Bacteroides* spp. populations (∼4 log_10_ cfu ml^-1^), and *Clostridium* spp. (∼1 log_10_ cfu ml^-1^). *Enterococcus* spp. populations recovered 4 days post antibiotic dosing and showed a further ∼1 log_10_ cfu ml^-1^ increase 11 days post cessation of antibiotic instillation. Total spores recovered to equilibration period levels (∼4 log_10_ cfu ml^-1^) approximately 4 days post completion of antibiotic dosing. *Bacteroides* spp. and *Clostridium* spp. populations recovered during rifaximin instillation. Interestingly, the *Bifidobacterium* spp. population increased ∼1.5 log_10_ cfu ml^-1^ and returned to equilibration period level approximately 5 days post antibiotic exposure.

Rifaximin κ induced declines in *Lactobacillus* spp. (∼1 log_10_ cfu ml^-1^), *Enterococcus* spp., total spores, and *Bifidobacterium* spp. (all ∼2 log_10_ cfu ml^-1^) and in *Bacteroides* spp. (∼3 log_10_ cfu ml^-1^) populations (Fig. 2c and Fig. S1c). All populations recovered before the end of antibiotic dosing, but *Bifidobacterium* spp. showed a further decline of ∼1 log_10_ cfu ml^-1^ approximately a week after antibiotic exposure ended. Following recovery, *Enterococcus* spp. and *Bacteroides* spp. populations remained ∼1 log_10_ cfu ml^-1^ higher compared with pre rifaximin κ instillation.

In all models, populations of total obligate anaerobes, total facultative anaerobes and lactose-fermenting Enterobacteriaceae remained stable throughout the experiment (Fig. S1). Whilst the levels of the total spores initially recovered to pre-antibiotic levels, in all models they later declined by ∼1 log10 cfu ml^-1^, which could be due to germination of these spores.

### Microbial diversity analysis by 16S sequencing in the gut models

The percentage of joined paired-end reads varied between 45.91% and 67.70%, and the number of reads per sample ranged from 19114 to 116233 (mean 74483) across all samples. Samples were normalised to the third lowest sample size, corresponding to 50736 reads, due to its considerable difference to the lowest values, 19114 and 25255. During equilibration phase, the global bacterial abundancies were similar in all models. Bifidobacteriaceae, Ruminococcaceae, Acidaminococcaceae, Lachnospiraceae and Rikenellaceae were the most abundant bacterial families, corresponding to >92% of the total relative abundance in all models (Fig. S2 and Table S1). This corresponded to 15 bacterial genera (*Oscillospira, Bifidobacterium, Megasphaera, Faecalibacterium, Coprococcus, Acidaminococcus, Ruminococcus, Sutterella, Bacteroides, Parabacteroides, Pyramidobacter, Clostridium, Dorea, Lactobacillus*, and *Lachnospira*) represented >99% of the overall relative abundance in all models (Fig. 3 and Table S2). In all models, genus *Oscillospira* and *Bifidobacterium* were highly abundant throughout the study. During rifaximin exposure, *Acidaminococcus* abundance increased in models A and B (20% and 11%, respectively). Instillation of rifaximin α and β also increased the relative abundance of *Lachnospira* in 1.7% and 3.3%, respectively, with this genus remaining highly prevalent up to the end of the experiment in both models. The relative abundance of genera *Oscillospira* declined (5.8% and 26.5%, with antibiotic dosing in models A and B, respectively. In models A and B, relative abundance of genus *Bacteroides* decreased at the start of antibiotic dosing but recovered still during antibiotic instillation (from 0.39% to 0.7% in model A; and from 0.16% to 1.19% in model B). These trends are consistent with those observed by direct enumeration (Fig. 2A and 2B). Differences between the effects of rifaximin α and β effects were also observed. During antibiotic dosing, genera *Megasphaera* and *Bifidobacterium* declined 8.5% and 5.3%, respectively, in model A but increased 3% and 10.7%, respectively, in model B. Relative abundance of genus *Faecalibacterium* increased 1.8% during rifaximin α instillation, and declined at the same level in presence of rifaximin β (Fig. 3 and Table S2).

**Figure 3.**
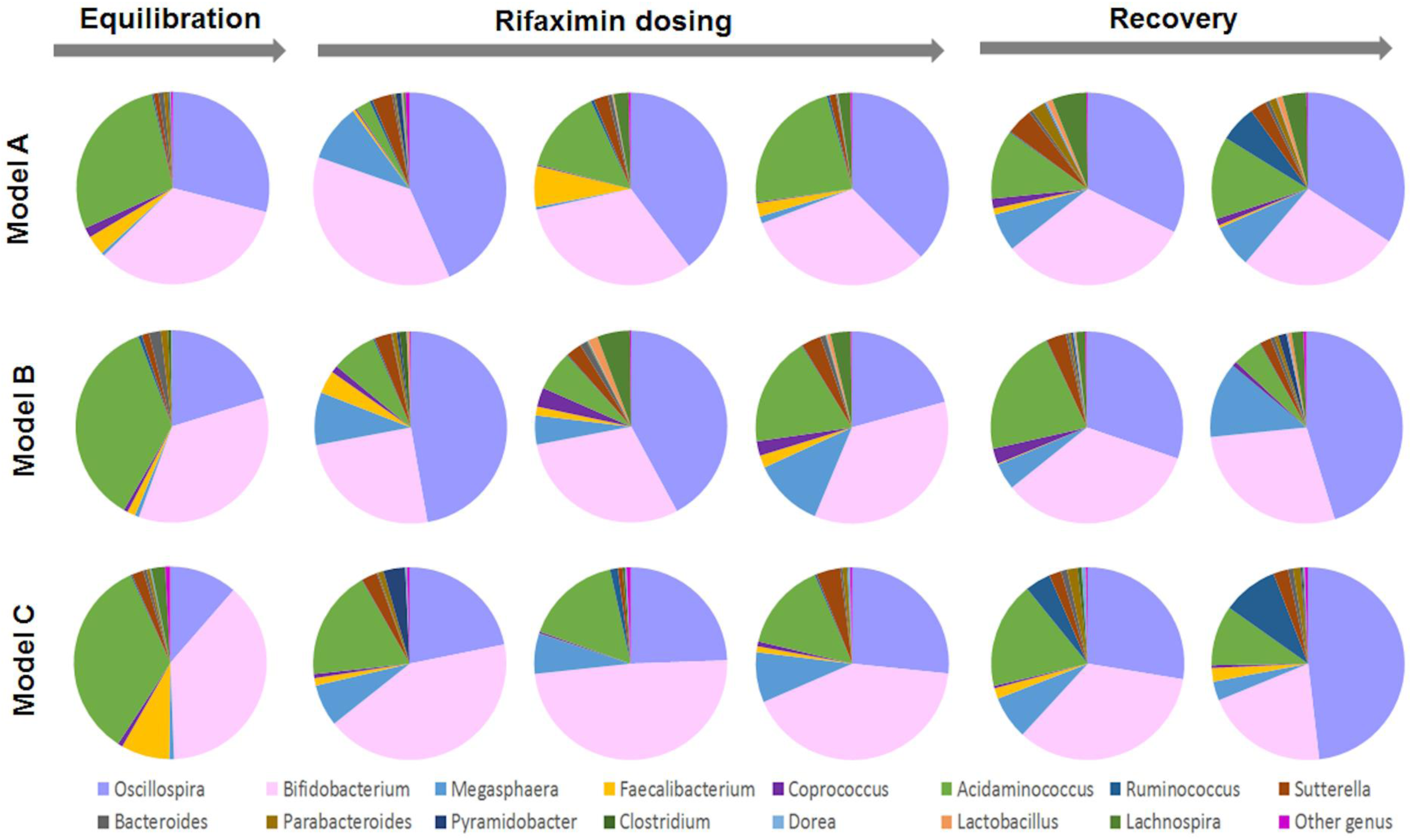
Pie charts constructed using bacterial OTUs assigned to a genus taxonomic level in models A, B and C throughout the experiment. Timeline for equilibration, rifaximin dosing and recovery periods are described in Figure 1. The legend shows the most abundant taxonomic genera.

In model C, genera *Bifidobacterium, Oscillospira*, and *Acidaminococcus* remained the most prevalent throughout the study. During rifaximin κ exposure, relative abundances of genera *Oscillospira, Megasphaera*, and *Sutterella*, increased 5.8%, 1.4% and 1.4%, respectively; whereas relative abundance of *Bifidobacterium* and *Acidaminococcus* declined 0.7% and 3.5%, respectively. In model C, sequencing data also showed a decline in genus *Bacteroides* from 0.22% to 0.015% between day 15 and day 20, with this genus recovering during antibiotic instillation (0.38% at day 23), which is consistent with the data observed by culture (Figure 2c). At the end of the experiment, models A and B showed similar relative abundance of *Bifidobacterium* (27% in A, 28.2% in B), whereas model C showed a slightly lower abundance of 20.62%. This is also consistent with the observations of bacterial culture, where at the end of the experiment, *Bifidobacterium* spp. counts are ∼1 log_10_ cfu ml^-1^ lower in model C, compared to models A and B (Fig. 3 and Table S2).

### Antimicrobial concentrations in Model A, B and C

Mean bioactive concentrations of the soluble fraction of rifaximin α, β and κ peaked at 43.1 mg l^-1^, 36.8 mg l^-1^ and 61.9 mg l^-1^ in vessel 3 of models A, B and C, respectively (Fig. 4a). Overall, the soluble phase of rifaximin was detected in vessel 3 from day 15 and persisted above the limit of detection (8 mg l^-1^) for the remainder of the experiment in all models. We detected sporadic increases in antibiotic concentrations in vessel 3 at day 31 and 35 in model A, at day 35 in model B, and at day 37 in model C. The insoluble phase of rifaximin α, β and κ peaked at 5400 mg l^-1^, 4635.5 mg l^-1^ and 4422.3 mg l^-1^ in vessel 3 of models A, B and C, respectively (Fig. 4b). Antibiotic concentrations in vessel 3 remained above the limit of detection for 25 days for rifaximin α and rifaximin κ, and until the end of the experiment for rifaximin β.

**Figure 4.**
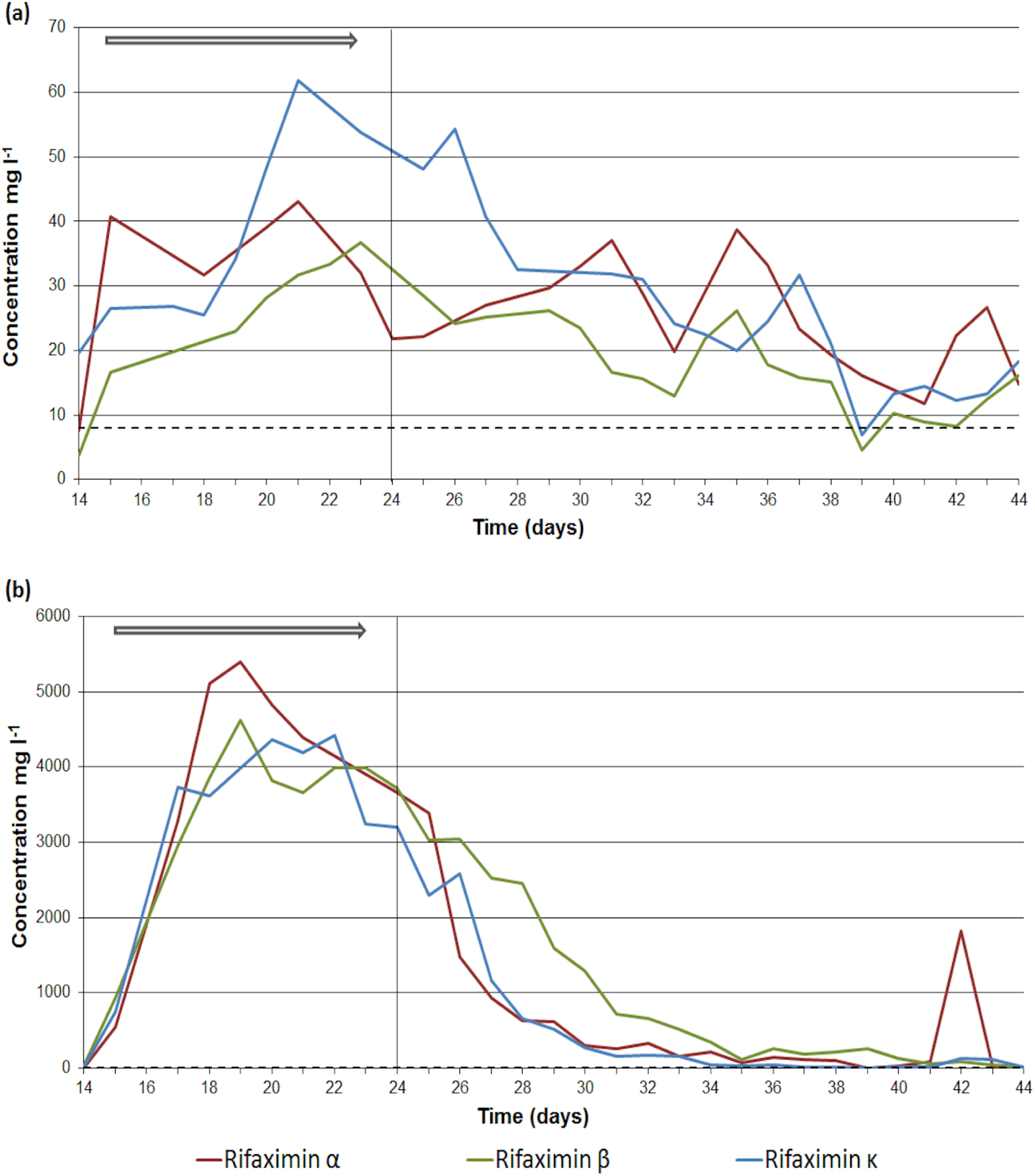
Antimicrobial concentration (mg l^-1^) in vessel 3 of model A, B and C regarding (a) soluble rifaximin, (b) insoluble phase of rifaximin. Horizontal arrow defines the period of antimicrobial instillation. Horizontal dotted line marks the limit of detection (8 mg l^-1^) for the antimicrobial bioassay.

## Discussion

Three *in vitro* gut models (A, B and C) were used to investigate the effects of three rifaximin formulations (α, β and κ) on the gut microbiome. Bacterial populations were monitored continuously by selective and non-selective culture throughout the study. Additionally, variations in global bacterial communities resulting from rifaximin instillation were assessed by 16S sequencing. All rifaximin formulations caused less bacterial disruption of gut microbiota populations in the chemostat models than observed with other antibiotics, [19-21, 24] which agrees with previous rifaximin studies [2, 11, 12, 15, 35]. Rifaximin formulations α, β and κ caused similar alterations in the gut microbiota, with some obligate (*Bacteroides* spp. and *Bifidobacterium* spp.) and facultative anaerobic (*Enterococcus* spp.) populations affected the most. Despite rifaximin low solubility, [2, 4] the soluble phase remained above 8 mg l^-1^ in all models throughout the experiment. The insoluble phase showed concentrations 100-fold higher than the soluble phase and similar to previously reported rifaximin’ faecal concentrations (8000 mg l^-1^) [17]. Biofilm formation in the gut model vessels was previously reported and hypothesised to have a role in fidaxomicin persistence in the gut model [21]. The occasional spikes in antibiotic concentration observed during recovery period could be associated with biofilm detachment from the vessel walls and subsequent release of antibiotic residue retained within the matrix. In all models, culture data showed a decrease of *Bacteroides* spp. populations at the start of rifaximin instillation, followed by a recovery to equilibration phase levels by the end of antibiotic dosing, and indeed before the elimination of bioactive antibiotic (which persisted in the insoluble phase for at least 3 weeks after dosing ended -Fig. 4). This is similar to the variations of the *Bacteroides* genus shown by the 16S sequencing data, particularly for models A and B. In the literature, *Bacteroides* abundance has been reported as unchanged [15] or increased [10, 12] after rifaximin dosing, however no testing was performed in those studies during antibiotic dosing. We observed that genus *Bacteroides* populations were affected by rifaximin but recovered within few days, showing similar results pre and post-antibiotic period. This could be associated with the expansion of sub-populations less susceptible to rifaximin, as MICs within this genus can show a wide susceptibility range, from 0.25 to >1024 mg l^-1^ [36]. This applies for instance to *B. fragilis*, a common component of the gut microbiota, present in these gut models (confirmed by MALDI-TOF identification, and whose polysaccharides are required for a normal immune system response and as such, may play a role in infection prevention [37]. Both bacterial culture and 16S sequencing data showed *Bifidobacterium* as highly prevalent in all models, with rifaximin exposure causing a decline in models A and C, and an increase in model B. Susceptibility of *Bifidobacterium* populations to each rifaximin formulation could explain these differences, as *Bifidobacterium* genus has been shown to be highly resistant to rifaximin, with MICs increasing up to 25 mg l^-1^ during antibiotic exposure [35, 36]. As observed in patient studies, [13, 15, 35] our culture and sequencing data showed recovery of the microbial populations affected by rifaximin instillation, although differences in the relative abundance of some bacterial groups were observed.

Overall, taxonomic analysis showed a variety of bacterial families and genera that otherwise would not have been evident. Rifaximin κ appeared to have the least disruptive effect on the colonic microbiota, with culture populations showing quicker recovery (i.e. during antibiotic dosing), and the sequencing data revealing the least variation in relative abundance of prevalent genera. Despite the proprietary nature of the formulations tested, this study contributes with novel data on the effects of rifamycins on the healthy human gut microbiota and supports the idea that antibiotic modification can be performed without compromise drug bioavailability or aggravating the effects to the intestinal microbiota.

## Supporting information

Table S1 - Abundance at Family level

Table S2 - Abundance at Genus level

## Funding

This study was initiated and financially supported by Teva Pharmaceuticals USA, Inc.. The Funder had no input on data analysis or in the preparation of the manuscript.

## Acknowledgements

The authors thank Miss Kate Owen and Mrs Sharie Shearman for the technical assistance.

## Ethical approval

The collection/use of faecal donations from healthy adult volunteers following informed consent was approved by the Leeds Institute of Health Sciences and Leeds Institute of Genetics, Health and Therapeutics and Leeds Institute of Molecular Medicine, University of Leeds joint ethics committee (reference HSLTLM/12/061).

---

**Supplementary Figure 1.**
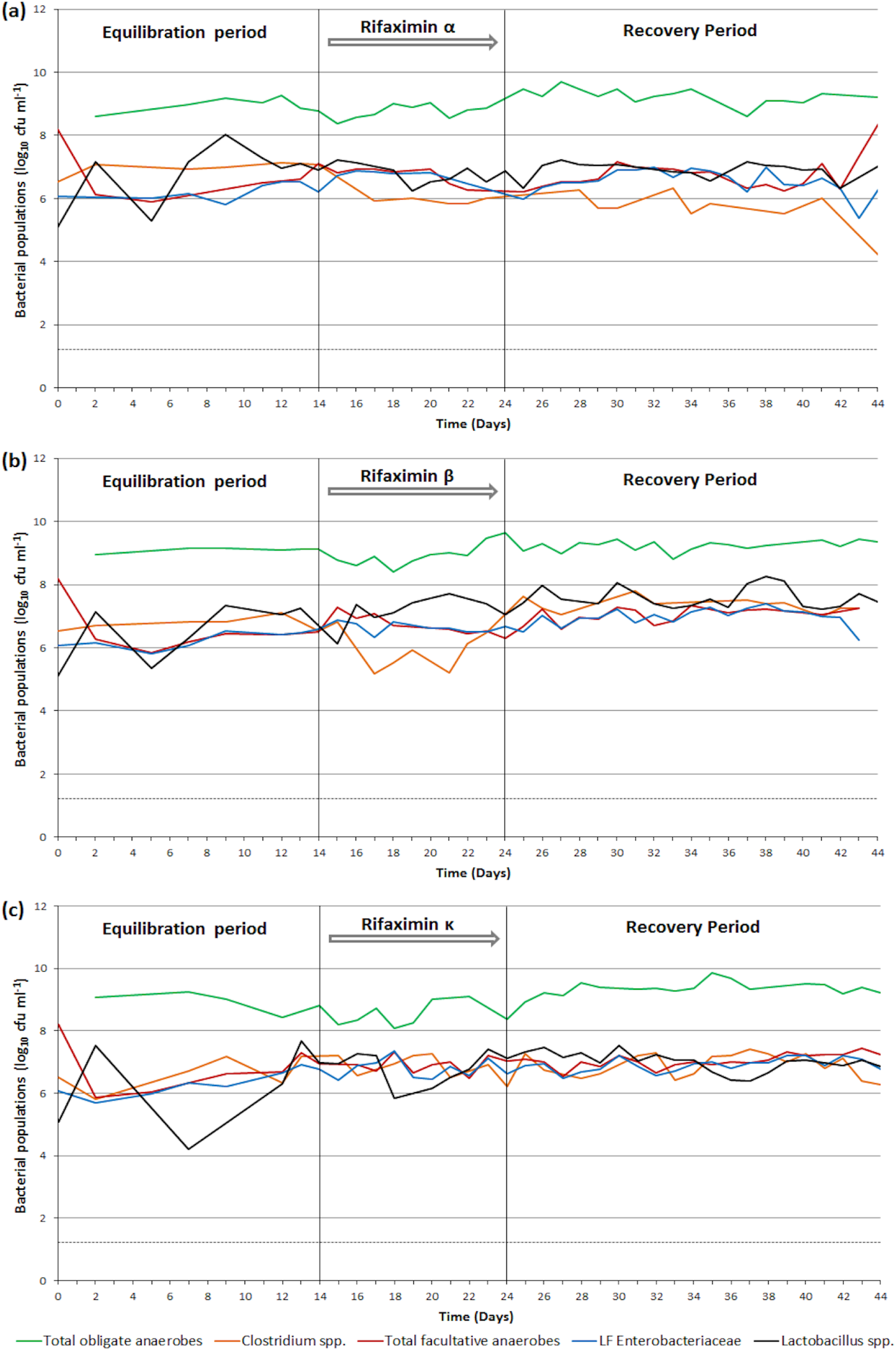
Mean gut microbiota populations (log_10_ cfu ml^-1^), in vessel 3 of (a) model A (rifaximin α dosing), (b) model B (rifaximin β dosing), (c) model C (rifaximin κ dosing). Horizontal dotted line marks the limit of detection for culture assay (∼1.2 log_10_ cfu ml^-1^). LF Enterobacteriaceae, lactose-fermenting Enterobacteriaceae.

**Supplementary Figure 2.**
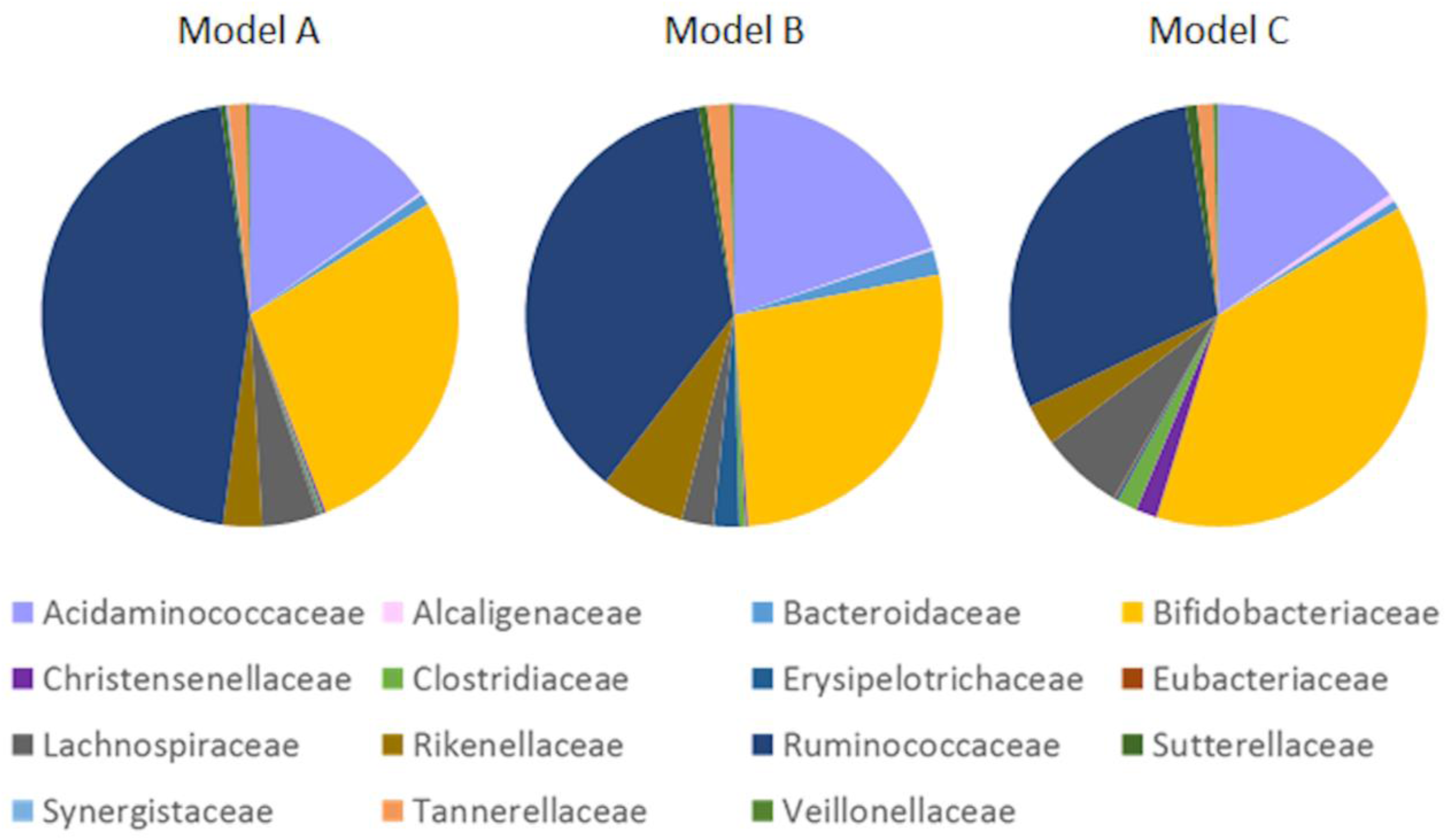
Pie charts constructed using bacterial OTUs assigned to a family taxonomic level in stationary phase of models A, B and C. The legend shows the most abundant taxonomic families.

**Supplementary Table 1.** Bacterial abundance at family level, in all models throughout the experiment.

**Supplementary Table 2.** Bacterial abundance at genus level, in all models throughout the experiment.

