## Supplementary material for "Profiling the effects of rifaximin on the healthy human colonic microbiota using a chemostat model": Table S1 - Abundance at Family level

| | Abundance % | | | | | | | | | | | | | | | | | |
|---|---|---|---|---|---|---|---|---|---|---|---|---|---|---|---|---|---|---|
| | Model A | | | | | | Model B | | | | | | Model C | | | | | |
| Bacterial Families | Stationary Phase | Rifaximin Period | | | Recovery Period | | Stationary Phase | Rifaximin Period | | | Recovery Period | | Stationary Phase | Rifaximin Period | | | Recovery Period | |
| | Day 5 | Day 15 | Day 20 | Day 23 | Day 33 | Day 44 | Day 5 | Day 15 | Day 20 | Day 23 | Day 33 | Day 44 | Day 5 | Day 15 | Day 20 | Day 23 | Day 33 | Day 44 |
| Ruminococcaceae | 45.5711 | 59.87955 | 53.37935 | 44.47407 | 35.15757 | 42.69115 | 36.60757 | 59.68607 | 52.99226 | 31.74623 | 35.74804 | 52.24818 | 29.47642 | 34.11489 | 30.77059 | 42.84358 | 34.35826 | 61.62512 |
| Bifidobacteriaceae | 27.85621 | 26.7232 | 29.38023 | 31.18263 | 31.33146 | 26.2666 | 26.99324 | 20.92946 | 25.23219 | 34.93402 | 32.69568 | 23.5731 | 37.99667 | 42.04853 | 38.56944 | 32.23034 | 32.61749 | 18.29241 |
| Lachnospiraceae | 4.327012 | 3.519152 | 4.213177 | 5.368818 | 10.4365 | 11.16767 | 2.416907 | 3.060841 | 9.301365 | 6.961728 | 8.331428 | 8.919614 | 6.33171 | 5.826574 | 3.807606 | 3.457313 | 6.37479 | 5.082999 |
| Acidaminococcaceae | 15.1375 | 0.988677 | 9.101266 | 15.68082 | 6.890805 | 8.394356 | 19.79295 | 4.158026 | 3.813528 | 12.56308 | 13.33232 | 2.528709 | 15.27813 | 8.013565 | 9.223656 | 9.434429 | 8.322217 | 4.60276 |
| Veillonellaceae | 0.308278 | 3.841002 | 0.304429 | 0.825913 | 3.819133 | 4.411502 | 0.401549 | 4.901363 | 2.80746 | 8.152839 | 2.777989 | 6.545164 | 0.3717 | 3.413865 | 5.341932 | 3.635321 | 3.399523 | 1.479167 |
| Tannerellaceae | 1.374655 | 0.545411 | 0.007659 | 0.016902 | 5.298086 | 2.505408 | 1.67661 | 1.473066 | 0.025416 | 0.197474 | 0.785001 | 1.18494 | 1.208024 | 0.01296 | 0.561959 | 1.441866 | 2.189079 | 1.569978 |
| Sutterellaceae | 0.441668 | 1.320164 | 1.530759 | 0.765218 | 2.643554 | 1.652081 | 0.676544 | 1.550187 | 1.513338 | 2.2349 | 2.049514 | 0.95755 | 0.894002 | 0.381238 | 2.500055 | 1.35385 | 0.951298 | 1.125737 |
| Lactobacillaceae | 0.002223 | 0.089617 | 0.192422 | 0.090658 | 0.758438 | 0.732274 | 0.006661 | 0.457538 | 2.005782 | 0.893333 | 0.231817 | 0.731303 | 0 | 0.00324 | 0.004425 | 0.025712 | 0.017495 | 0.010636 |
| Bacteroidaceae | 0.889264 | 0.24476 | 0.596412 | 0.1575 | 0.475021 | 0.549701 | 1.897367 | 0.121837 | 0.880045 | 0.783625 | 0.389325 | 0.452494 | 0.666496 | 0.02052 | 0.316379 | 0.159218 | 0.743543 | 0.590685 |
| Alcaligenaceae | 0.16896 | 0.637919 | 0.438454 | 0.199756 | 0.882183 | 0.521919 | 0.176035 | 0.550861 | 0.514683 | 0.529731 | 0.661154 | 0.348512 | 0.551141 | 0.180359 | 0.825239 | 0.418319 | 0.5172 | 0.652862 |
| Eubacteriaceae | 0.068177 | 0.091544 | 0.019146 | 0.365707 | 1.045846 | 0.450477 | 0.038062 | 0.105636 | 0.02118 | 0.137918 | 0.362015 | 0.353082 | 0.102538 | 0.186839 | 0.03208 | 0.126584 | 0.22853 | 0.229074 |
| Erysipelotrichaceae | 0.132649 | 0.119489 | 0.156044 | 0.218195 | 0.187614 | 0.22127 | 1.972539 | 0.090082 | 0.109079 | 0.12538 | 0.238168 | 0.46392 | 0.256345 | 1.349994 | 0.396026 | 0.358983 | 0.262427 | 0.439332 |
| Clostridiaceae | 0.210459 | 0.158998 | 0.186678 | 0.464816 | 0.243498 | 0.171658 | 0.475769 | 0.826291 | 0.308175 | 0.316585 | 0.178467 | 0.137119 | 1.54768 | 4.287581 | 1.484546 | 0.177019 | 9.830078 | 3.342033 |
| Desulfovibrionaceae | 0.012598 | 0.160925 | 0.191465 | 0.126768 | 0.073848 | 0.127007 | 0.023788 | 0.040828 | 0.027534 | 0.028211 | 0.038107 | 0.114266 | 0.02243 | 0.00324 | 0.016593 | 0.015823 | 0.007654 | 0.102265 |
| Enterobacteriaceae | 0.002964 | 0.015418 | 0.018189 | 0.006915 | 0.024949 | 0.036713 | 0.002855 | 0.01685 | 0.018003 | 0.018807 | 0.023499 | 0.202251 | 0.006409 | 0.01188 | 0.018806 | 0.026701 | 0.017495 | 0.080176 |
| Coriobacteriaceae | 0.001482 | 0.000964 | 0.031592 | 0 | 0.049897 | 0.021829 | 0.032352 | 0 | 0.001059 | 0 | 0.031121 | 0.020568 | 0.006409 | 0.0162 | 0.021018 | 0.005934 | 0.019682 | 0.015544 |
| Synergistaceae | 0.152657 | 0.516502 | 0.002872 | 0 | 0.185618 | 0.014884 | 0.001903 | 0.330516 | 0.016944 | 0.015673 | 0.292788 | 1.028395 | 0.009613 | 0.00216 | 0.006637 | 2.470357 | 0 | 0.175078 |
| Comamonadaceae | 0.003705 | 0.014454 | 0.01436 | 0.005378 | 0.026944 | 0.013891 | 0.006661 | 0.006481 | 0.011649 | 0.021942 | 0.016513 | 0.009141 | 0.009613 | 0.00108 | 0.025443 | 0.015823 | 0.010934 | 0.006545 |
| Eggerthellaceae | 0.002964 | 0.001927 | 0 | 0.006915 | 0.013971 | 0.013891 | 0.000952 | 0.00324 | 0 | 0.009404 | 0.007621 | 0.012569 | 0.006409 | 0.00108 | 0.005531 | 0.011867 | 0.005467 | 0.002454 |
| Rikenellaceae | 3.006454 | 0.048181 | 0.016274 | 0.009219 | 0.020957 | 0.00893 | 6.488539 | 1.00127 | 0.022239 | 0.021942 | 0.042553 | 0.030852 | 3.146629 | 0.05508 | 5.978008 | 1.672287 | 0.030616 | 0.431151 |
| Christensenellaceae | 0.226762 | 1.052277 | 0.192422 | 0.009988 | 0.401174 | 0.005953 | 0.18555 | 0.666865 | 0.34524 | 0.278971 | 1.744024 | 0.041136 | 1.647014 | 0.00216 | 0.022124 | 0.050436 | 0.03171 | 0.041724 |
| Helicobacteraceae | 0.004446 | 0.006745 | 0.005744 | 0.002305 | 0.009979 | 0.005953 | 0.000952 | 0.001944 | 0.005295 | 0.009404 | 0.003176 | 0.007999 | 0.012817 | 0.0054 | 0.005531 | 0.003956 | 0.002187 | 0.000818 |
| Peptostreptococcaceae | 0.031865 | 0 | 0.000957 | 0.002305 | 0.002994 | 0.002977 | 0.035207 | 0.001944 | 0 | 0.003135 | 0.000635 | 0.002285 | 0.400538 | 0.00216 | 0.004425 | 0.001978 | 0.001093 | 0.003272 |
| Prevotellaceae | 0.005187 | 0.002891 | 0 | 0.001537 | 0.002994 | 0.001984 | 0.027595 | 0.00324 | 0.002118 | 0 | 0.000635 | 0 | 0.003204 | 0 | 0.003319 | 0 | 0.006561 | 0.002454 |
| Verrucomicrobiaceae | 0.002964 | 0.001927 | 0.001915 | 0.001537 | 0.001996 | 0.001984 | 0.002855 | 0.001944 | 0.002118 | 0 | 0 | 0.001143 | 0 | 0.00216 | 0 | 0.001978 | 0 | 0 |
| Oxalobacteraceae | 0.000741 | 0 | 0.002872 | 0.001537 | 0.005988 | 0.001984 | 0.003806 | 0.003888 | 0.001059 | 0 | 0.006986 | 0.002285 | 0.009613 | 0 | 0.006637 | 0.003956 | 0.001093 | 0 |
| Odoribacteraceae | 0.008152 | 0 | 0 | 0.001537 | 0.000998 | 0.000992 | 0.010467 | 0.000648 | 0 | 0.003135 | 0 | 0 | 0.006409 | 0 | 0 | 0 | 0 | 0.000818 |
| Deferribacteraceae | 0.000741 | 0.003854 | 0.001915 | 0.001537 | 0.001996 | 0.000992 | 0.001903 | 0 | 0.004236 | 0.003135 | 0.003176 | 0.002285 | 0 | 0.00432 | 0 | 0.004945 | 0.002187 | 0.001636 |
| Methanobacteriaceae | 0 | 0.002891 | 0 | 0.000768 | 0.001996 | 0.000992 | 0 | 0.000648 | 0.002118 | 0 | 0.000635 | 0 | 0 | 0 | 0 | 0 | 0 | 0 |
| Bacillaceae | 0 | 0.000964 | 0.000957 | 0.000768 | 0 | 0.000992 | 0.001903 | 0 | 0.003177 | 0 | 0.00127 | 0.065132 | 0 | 0.00432 | 0.003319 | 0 | 0.002187 | 0.006545 |
| Anaeroplasmataceae | 0 | 0 | 0.001915 | 0 | 0 | 0.000992 | 0 | 0 | 0 | 0.003135 | 0 | 0 | 0 | 0 | 0 | 0.000989 | 0.001093 | 0.000818 |
| Enterococcaceae | 0 | 0 | 0 | 0 | 0 | 0.000992 | 0.002855 | 0 | 0 | 0 | 0.000635 | 0.002285 | 0 | 0.02916 | 0.042036 | 0 | 0.029523 | 0.068722 |
| Victivallaceae | 0.043722 | 0 | 0 | 0 | 0.000998 | 0 | 0.027595 | 0.000648 | 0.002118 | 0 | 0.00127 | 0 | 0.006409 | 0 | 0 | 0 | 0 | 0 |
| Peptococcaceae | 0.001482 | 0.006745 | 0.006701 | 0.006915 | 0.000998 | 0 | 0.004758 | 0.003888 | 0.003177 | 0.006269 | 0.00127 | 0.007999 | 0.009613 | 0.0108 | 0 | 0.015823 | 0.005467 | 0.002454 |
| Corynebacteriaceae | 0.001482 | 0 | 0 | 0 | 0 | 0 | 0 | 0 | 0 | 0 | 0 | 0 | 0 | 0.00216 | 0 | 0.031646 | 0 | 0 |
| Gracilibacteraceae | 0.000741 | 0.002891 | 0 | 0.000768 | 0 | 0 | 0.001903 | 0.001296 | 0.001059 | 0 | 0.000635 | 0.001143 | 0 | 0.00108 | 0.001106 | 0.001978 | 0.001093 | 0.000818 |
| Rhodocyclaceae | 0.000741 | 0 | 0 | 0 | 0 | 0 | 0 | 0 | 0.001059 | 0 | 0 | 0 | 0 | 0 | 0 | 0 | 0.001093 | 0 |
| Micrococcaceae | 0 | 0.000964 | 0 | 0 | 0 | 0 | 0 | 0 | 0 | 0 | 0 | 0 | 0 | 0 | 0 | 0 | 0.008748 | 0.009817 |
| Burkholderiaceae | 0 | 0 | 0.000957 | 0 | 0 | 0 | 0 | 0 | 0.002118 | 0 | 0.000635 | 0 | 0 | 0 | 0.001106 | 0 | 0.001093 | 0 |
| Pasteurellaceae | 0 | 0 | 0.000957 | 0 | 0 | 0 | 0.000952 | 0 | 0 | 0 | 0 | 0 | 0 | 0.00108 | 0.001106 | 0 | 0 | 0 |
| Moraxellaceae | 0 | 0 | 0.000957 | 0 | 0 | 0 | 0 | 0 | 0 | 0 | 0 | 0 | 0 | 0 | 0 | 0 | 0 | 0 |
| Brachyspiraceae | 0 | 0 | 0.000957 | 0 | 0 | 0 | 0 | 0 | 0 | 0 | 0 | 0 | 0 | 0 | 0 | 0 | 0 | 0 |
| Streptococcaceae | 0 | 0 | 0 | 0.000768 | 0.000998 | 0 | 0.001903 | 0.000648 | 0.001059 | 0 | 0.000635 | 0 | 0 | 0 | 0 | 0 | 0 | 0 |
| Thermoactinomycetaceae | 0 | 0 | 0 | 0.000768 | 0 | 0 | 0 | 0 | 0 | 0 | 0 | 0 | 0 | 0 | 0 | 0 | 0 | 0 |
| Selenomonadaceae | 0 | 0 | 0 | 0.000768 | 0 | 0 | 0.000952 | 0 | 0 | 0 | 0 | 0 | 0 | 0.00108 | 0 | 0 | 0 | 0 |
| Aerococcaceae | 0 | 0 | 0 | 0 | 0.000998 | 0 | 0 | 0 | 0.001059 | 0 | 0 | 0 | 0 | 0 | 0 | 0 | 0 | 0 |
| Carnobacteriaceae | 0 | 0 | 0 | 0 | 0 | 0 | 0 | 0 | 0 | 0 | 0 | 0 | 0 | 0 | 0 | 0 | 0 | 0.001636 |
| Planococcaceae | 0 | 0 | 0 | 0 | 0 | 0 | 0 | 0.000648 | 0 | 0 | 0 | 0.002285 | 0 | 0 | 0 | 0 | 0 | 0 |
| Actinomycetaceae | 0 | 0 | 0 | 0 | 0 | 0 | 0 | 0 | 0 | 0 | 0.00127 | 0.001143 | 0 | 0 | 0.001106 | 0 | 0 | 0 |
| Akkermansiaceae | 0 | 0 | 0 | 0 | 0 | 0 | 0 | 0 | 0.001059 | 0 | 0 | 0 | 0 | 0.00324 | 0.002212 | 0 | 0 | 0 |
| Peptoniphilaceae | 0 | 0 | 0 | 0 | 0 | 0 | 0 | 0 | 0 | 0 | 0 | 0 | 0 | 0 | 0 | 0 | 0 | 0.001636 |
| Caldicoprobacteraceae | 0 | 0 | 0 | 0 | 0 | 0 | 0 | 0.000648 | 0 | 0 | 0 | 0 | 0 | 0 | 0 | 0 | 0.001093 | 0.000818 |
| Clostridiales Family XIII. Incertae Sedis | 0 | 0 | 0 | 0 | 0 | 0 | 0 | 0 | 0 | 0 | 0 | 0 | 0.016022 | 0 | 0 | 0 | 0 | 0 |
| Brevibacteriaceae | 0 | 0 | 0 | 0 | 0 | 0 | 0 | 0 | 0 | 0 | 0 | 0.001143 | 0 | 0 | 0 | 0 | 0 | 0 |
| Staphylococcaceae | 0 | 0 | 0 | 0 | 0 | 0 | 0 | 0 | 0 | 0 | 0 | 0 | 0 | 0 | 0 | 0.000989 | 0 | 0 |
| Campylobacteraceae | 0 | 0 | 0 | 0 | 0 | 0 | 0 | 0.000648 | 0 | 0 | 0 | 0 | 0 | 0 | 0 | 0 | 0 | 0 |
