## Supplementary material for "Profiling the effects of rifaximin on the healthy human colonic microbiota using a chemostat model": Table S2 - Abundance at Genus level

**Model A**

| Bacterial Genus | Stationary Phase | Rifaximin Period | | | Recovery Period | |
|---|---|---|---|---|---|---|
| | Day 5 | Day 15 | Day 20 | Day 23 | Day 33 | Day 44 |
| Oscillospira | 29.019716 | 43.289219 | 39.73106 | 37.450863 | 32.395287 | 34.198063 |
| Bifidobacterium | 33.633713 | 37.042725 | 31.787823 | 31.739561 | 31.943886 | 27.00218 |
| Megasphaera | 0.565121 | 9.663425 | 0.470049 | 1.182694 | 6.364948 | 7.199161 |
| Faecalibacterium | 3.275479 | 0.414321 | 6.869264 | 2.244087 | 1.037052 | 0.41625 |
| Coprococcus | 1.675923 | 0.255904 | 0.22015 | 0.20217 | 1.64955 | 1.150424 |
| Acidaminococcus | 28.29353 | 2.4908 | 14.14164 | 22.919334 | 11.587126 | 13.860802 |
| Ruminococcus | 0.290197 | 0.533743 | 0.553349 | 0.403217 | 0.192979 | 6.170007 |
| Sutterella | 0.827548 | 3.338939 | 2.378509 | 1.118674 | 4.445228 | 2.728569 |
| Bacteroides | 0.89975 | 0.389949 | 0.708049 | 0.17746 | 0.599074 | 0.680094 |
| Parabacteroides | 0.715079 | 0.25103 | 0 | 0.010108 | 2.265405 | 1.202865 |
| Pyramidobacter | 0.206887 | 0.870074 | 0.002975 | 0 | 0.199691 | 0.014749 |
| Clostridium | 0.131908 | 0.129171 | 0.014875 | 0.015724 | 0.010068 | 0.01311 |
| Dorea | 0.126354 | 0.148668 | 0.07735 | 0.080868 | 0.342328 | 0.12127 |
| Lactobacillus | 0 | 0.114548 | 0.2023 | 0.069636 | 0.939723 | 0.920994 |
| Blautia | 0.026382 | 0.17304 | 0.055037 | 0.055035 | 0.010068 | 0.011471 |
| Desulfovibrio | 0.012497 | 0.282713 | 0.229075 | 0.150504 | 0.063767 | 0.145851 |
| Christensenella | 0.002777 | 0.026809 | 0 | 0 | 0 | 0 |
| Lachnospira | 0.031936 | 0.355829 | 2.469246 | 2.087966 | 5.791046 | 4.051064 |
| Phascolarctobacterium | 0.069425 | 0.009749 | 0 | 0.004493 | 0 | 0.003278 |
| Pediococcus | 0 | 0.004874 | 0.004462 | 0.001123 | 0 | 0 |
| Holdemania | 0.020828 | 0.124296 | 0.02975 | 0.06402 | 0.031883 | 0.021304 |
| Anaerotruncus | 0.022216 | 0.012186 | 0 | 0.001123 | 0.001678 | 0 |
| Anaerofustis | 0.062483 | 0.019497 | 0 | 0 | 0 | 0 |
| Eggerthella | 0.004166 | 0.004874 | 0 | 0.010108 | 0.023493 | 0.022943 |
| Turicibacter | 0.051375 | 0.01706 | 0.001487 | 0.005616 | 0.005034 | 0.006555 |
| Bilophila | 0.002777 | 0.002437 | 0.001487 | 0.002246 | 0.010068 | 0.014749 |
| Prevotella | 0.001389 | 0.002437 | 0 | 0 | 0.001678 | 0 |
| Peptococcus | 0.001389 | 0.002437 | 0 | 0 | 0 | 0 |
| Allobaculum | 0.001389 | 0.002437 | 0.001487 | 0 | 0 | 0 |
| Odoribacter | 0.00972 | 0 | 0 | 0 | 0 | 0.001639 |
| Corynebacterium | 0.002777 | 0 | 0 | 0 | 0 | 0 |
| Collinsella | 0.002777 | 0 | 0.049087 | 0 | 0.083904 | 0.036053 |
| Coprobacillus | 0.002777 | 0 | 0 | 0 | 0 | 0 |
| Butyricimonas | 0.001389 | 0 | 0 | 0 | 0 | 0 |
| Flexispira | 0.001389 | 0 | 0 | 0.001123 | 0 | 0.001639 |
| Escherichia | 0.001389 | 0 | 0 | 0 | 0 | 0 |
| Slackia | 0.001389 | 0 | 0 | 0 | 0 | 0 |
| Dehalobacterium | 0.001389 | 0.004874 | 0 | 0 | 0 | 0 |
| Anaerostipes | 0.001389 | 0 | 0 | 0 | 0 | 0 |
| Veillonella | 0.001389 | 0.002437 | 0 | 0 | 0 | 0 |
| Mucispirillum | 0 | 0.002437 | 0 | 0 | 0 | 0 |
| Acinetobacter | 0 | 0 | 0.001487 | 0 | 0 | 0 |
| Micrococcus | 0 | 0.002437 | 0 | 0 | 0 | 0 |
| Enterococcus | 0 | 0 | 0 | 0 | 0 | 0.001639 |
| Oxobacter | 0 | 0.004874 | 0 | 0 | 0 | 0 |
| Roseburia | 0 | 0 | 0 | 0 | 0.001678 | 0 |
| Anaerofilum | 0 | 0.002437 | 0 | 0 | 0 | 0 |
| Megamonas | 0 | 0 | 0 | 0.001123 | 0 | 0 |
| Anaeroplasma | 0 | 0 | 0 | 0 | 0 | 0.001639 |
| Methanobrevibacter | 0 | 0.007312 | 0 | 0.001123 | 0.003356 | 0.001639 |

Abundance %

**Model B**

| Bacterial Genus | Stationary Phase | Rifaximin Period | | | Recovery Period | |
|---|---|---|---|---|---|---|
| | Day 5 | Day 15 | Day 20 | Day 23 | Day 33 | Day 44 |
| Oscillospira | 20.113325 | 47.22475 | 42.10556 | 20.71148 | 30.27105 | 45.24741 |
| Bifidobacterium | 35.095963 | 24.81319 | 29.85887 | 35.59251 | 33.94846 | 28.16672 |
| Megasphaera | 0.711731 | 8.899494 | 4.906878 | 11.87341 | 4.472561 | 12.70487 |
| Faecalibacterium | 1.323268 | 3.839276 | 1.476538 | 2.069762 | 0.146068 | 0 |
| Coprococcus | 0.701366 | 1.225679 | 3.26628 | 2.453223 | 2.606594 | 0.822843 |
| Acidaminococcus | 35.74205 | 7.592103 | 6.70967 | 18.51698 | 21.59132 | 4.946001 |
| Ruminococcus | 0.606354 | 0.266452 | 0.149145 | 0.14322 | 0.116237 | 0.138631 |
| Sutterella | 1.228255 | 2.83268 | 2.664106 | 3.294063 | 3.319447 | 1.873756 |
| Bacteroides | 1.983174 | 0.16224 | 1.187569 | 0.887041 | 0.485522 | 0.699864 |
| Parabacteroides | 1.153972 | 0.795803 | 0.005593 | 0.0693 | 0.385743 | 0.641728 |
| Pyramidobacter | 0.003455 | 0.460666 | 0.020507 | 0.01386 | 0.36517 | 1.406435 |
| Clostridium | 0.528616 | 1.187783 | 0.160331 | 0.02772 | 0.02263 | 0.004472 |
| Dorea | 0.120925 | 0.099475 | 0.139824 | 0.1617 | 0.27362 | 0.263846 |
| Lactobacillus | 0.005183 | 0.348164 | 1.63314 | 0.586741 | 0.169727 | 0.565704 |
| Lachnospira | 0.022458 | 0.05329 | 5.484815 | 3.400323 | 1.59749 | 2.023567 |
| Anaerotruncus | 0.010365 | 0.024869 | 0.001864 | 0 | 0.003086 | 0 |
| Holdemania | 0.031095 | 0.005921 | 0.007457 | 0.00462 | 0.010286 | 0.071552 |
| Prevotella | 0.036278 | 0.001184 | 0 | 0 | 0 | 0 |
| Blautia | 0.081193 | 0.045001 | 0.137959 | 0.09702 | 0.054518 | 0.013416 |
| Phascolarctobacterium | 0.191753 | 0.005921 | 0.003729 | 0 | 0.002057 | 0.002236 |
| Bilophila | 0.005183 | 0.013027 | 0.022372 | 0.01386 | 0.002057 | 0.008944 |
| Anaerofustis | 0.015548 | 0.004737 | 0.001864 | 0.00924 | 0.011315 | 0 |
| Christensenella | 0 | 0.001184 | 0.001864 | 0 | 0.013372 | 0.038012 |
| Desulfovibrio | 0.031095 | 0.047369 | 0.016779 | 0.0231 | 0.051432 | 0.163227 |
| Eggerthella | 0 | 0.005921 | 0 | 0.01386 | 0.012344 | 0.024596 |
| Turicibacter | 0.171023 | 0.010658 | 0.007457 | 0.00462 | 0.006172 | 0.01118 |
| Collinsella | 0.051825 | 0 | 0.001864 | 0 | 0.050404 | 0.040248 |
| Butyricimonas | 0.01382 | 0 | 0 | 0 | 0 | 0 |
| Alistipes | 0.005183 | 0 | 0 | 0 | 0 | 0 |
| Dehalobacterium | 0.005183 | 0.001184 | 0 | 0.00462 | 0 | 0 |
| Enterococcus | 0.003455 | 0 | 0 | 0 | 0 | 0 |
| Slackia | 0.001728 | 0 | 0 | 0 | 0 | 0 |
| Peptococcus | 0.001728 | 0.001184 | 0.001864 | 0 | 0.001029 | 0.002236 |
| Coprobacillus | 0.001728 | 0 | 0 | 0 | 0 | 0 |
| Megamonas | 0.001728 | 0 | 0 | 0 | 0 | 0 |
| Mucispirillum | 0 | 0 | 0.001864 | 0.00462 | 0.001029 | 0 |
| Odoribacter | 0 | 0 | 0 | 0.00462 | 0 | 0 |
| Lautropia | 0 | 0 | 0 | 0 | 0.001029 | 0 |
| Flexispira | 0 | 0 | 0 | 0 | 0 | 0.002236 |
| Citrobacter | 0 | 0.001184 | 0 | 0 | 0.008229 | 0.053664 |
| Akkermansia | 0 | 0 | 0.001864 | 0 | 0 | 0 |
| Brevibacterium | 0 | 0 | 0 | 0 | 0 | 0.002236 |
| Anaerobacillus | 0 | 0 | 0.001864 | 0 | 0 | 0 |
| Bacillus | 0 | 0 | 0 | 0 | 0 | 0.049192 |
| Pediococcus | 0 | 0.021316 | 0.01305 | 0 | 0 | 0 |
| Streptococcus | 0 | 0 | 0 | 0 | 0 | 0 |
| Caldicoprobacter | 0 | 0.001184 | 0 | 0 | 0 | 0 |
| Anaerostipes | 0 | 0.001184 | 0.001864 | 0 | 0 | 0.004472 |
| Allobaculum | 0 | 0.002368 | 0.003729 | 0 | 0 | 0 |
| Catenibacterium | 0 | 0 | 0 | 0 | 0 | 0 |
| Dialister | 0 | 0.002368 | 0 | 0.00924 | 0 | 0 |
| Veillonella | 0 | 0.001184 | 0 | 0.00924 | 0 | 0.006708 |
| Methanobrevibacter | 0 | 0 | 0.001864 | 0 | 0 | 0 |

Abundance %

**Model C**

| Bacterial Genus | Stationary Phase | Rifaximin Period | | | Recovery Period | |
|---|---|---|---|---|---|---|
| | Day 5 | Day 15 | Day 20 | Day 23 | Day 33 | Day 44 |
| Oscillospira | 11.373931 | 21.8063 | 24.45204 | 26.64747 | 27.59276 | 48.16664 |
| Bifidobacterium | 38.018551 | 42.49017 | 48.83443 | 41.83551 | 34.22986 | 20.6206 |
| Megasphaera | 0.790855 | 7.043123 | 6.777964 | 8.433627 | 7.26607 | 3.205025 |
| Faecalibacterium | 8.109857 | 1.245854 | 0.032649 | 0.976412 | 1.783661 | 2.270897 |
| Coprococcus | 0.848372 | 0.709712 | 0.319962 | 0.83156 | 0.441221 | 0.499275 |
| Acidaminococcus | 34.093033 | 18.39466 | 16.14392 | 14.90907 | 17.86008 | 10.05351 |
| Ruminococcus | 0.215688 | 0.111857 | 1.292906 | 0.323682 | 4.301908 | 9.346647 |
| Sutterella | 2.005895 | 2.640207 | 0.768343 | 4.04156 | 2.041822 | 2.462375 |
| Bacteroides | 0.567978 | 0.217928 | 0.015236 | 0.384485 | 1.030299 | 0.878653 |
| Parabacteroides | 0.496082 | 0.939212 | 0.00653 | 0.692072 | 1.753151 | 1.249083 |
| Pyramidobacter | 0.00719 | 3.654632 | 0 | 0.007153 | 0 | 0.238006 |
| Clostridium | 0.107844 | 0.017357 | 0.393966 | 0.162735 | 0.692342 | 0.245164 |
| Dorea | 0.165361 | 0.244928 | 0.25684 | 0.364813 | 0.675914 | 0.189689 |
| Lactobacillus | 0 | 0.025071 | 0.002177 | 0.003577 | 0.011735 | 0.014316 |
| Lachnospira | 2.343806 | 0.017357 | 0.017413 | 0.012518 | 0.004694 | 0.010737 |
| Christensenella | 0.00719 | 0.028928 | 0 | 0 | 0 | 0 |
| Blautia | 0.273204 | 0.057857 | 0.037002 | 0.02146 | 0.086836 | 0.044738 |
| Corynebacterium | 0 | 0.061714 | 0.004353 | 0 | 0 | 0 |
| Holdemania | 0.028758 | 0.167785 | 0.185012 | 0.062591 | 0.028163 | 0.143161 |
| Desulfotomaculum | 0 | 0.023143 | 0 | 0 | 0 | 0 |
| Bilophila | 0 | 0.0135 | 0.002177 | 0.014306 | 0.004694 | 0.00179 |
| Anaerotruncus | 0.021569 | 0.001929 | 0 | 0 | 0 | 0 |
| Eggerthella | 0.014379 | 0.023143 | 0.002177 | 0.008942 | 0.011735 | 0.005369 |
| Anaerofustis | 0.021569 | 0.0135 | 0.017413 | 0.003577 | 0.049285 | 0.005369 |
| Veillonella | 0 | 0.023143 | 0.052239 | 0.078685 | 0 | 0.00179 |
| Turicibacter | 0.150981 | 0.001929 | 0.004353 | 0.062591 | 0.002347 | 0.010737 |
| Anaerovorax | 0.00719 | 0 | 0 | 0 | 0 | 0 |
| Phascolarctobacterium | 0.186929 | 0.003857 | 0.00653 | 0.001788 | 0.002347 | 0.014316 |
| Staphylococcus | 0 | 0.001929 | 0 | 0 | 0 | 0 |
| Desulfovibrio | 0.028758 | 0.001929 | 0.002177 | 0.007153 | 0.002347 | 0.141372 |
| Collinsella | 0.014379 | 0.009643 | 0.032649 | 0.033978 | 0.042245 | 0.034001 |
| Mucispirillum | 0 | 0.003857 | 0 | 0 | 0.002347 | 0 |
| Megamonas | 0 | 0 | 0.002177 | 0 | 0 | 0 |
| Helicobacter | 0.00719 | 0 | 0 | 0 | 0 | 0 |
| Prevotella | 0 | 0 | 0 | 0 | 0 | 0.00179 |
| Pediococcus | 0 | 0.001929 | 0 | 0 | 0 | 0 |
| Enterococcus | 0 | 0 | 0.032649 | 0.041131 | 0.037551 | 0.087686 |
| Odoribacter | 0.00719 | 0 | 0 | 0 | 0 | 0.00179 |
| Bulleidia | 0 | 0 | 0.280783 | 0.026825 | 0.011735 | 0.008948 |
| Oxobacter | 0 | 0 | 0 | 0 | 0.004694 | 0 |
| Anaeroplasma | 0 | 0 | 0 | 0 | 0 | 0.00179 |
| Akkermansia | 0 | 0 | 0.00653 | 0.003577 | 0 | 0 |
| Anaerobacillus | 0 | 0 | 0.00653 | 0.003577 | 0 | 0.007158 |
| Peptococcus | 0 | 0 | 0.00653 | 0 | 0.002347 | 0 |
| Haemophilus | 0 | 0 | 0.002177 | 0.001788 | 0 | 0 |
| Gracilibacter | 0 | 0 | 0.002177 | 0 | 0 | 0.00179 |
| Allobaculum | 0.064706 | 0 | 0 | 0 | 0 | 0 |
| Butyricimonas | 0.00719 | 0 | 0 | 0 | 0 | 0 |
| Roseburia | 0.00719 | 0 | 0 | 0 | 0 | 0 |
| Dialister | 0.00719 | 0 | 0 | 0 | 0 | 0 |
| Lautropia | 0 | 0 | 0 | 0.001788 | 0 | 0 |
| Flexispira | 0 | 0 | 0 | 0 | 0 | 0.00179 |
| Citrobacter | 0 | 0 | 0 | 0 | 0.002347 | 0.007158 |
| Micrococcus | 0 | 0 | 0 | 0 | 0.018775 | 0.021474 |
| Caldicoprobacter | 0 | 0 | 0 | 0 | 0.002347 | 0 |
| Acetobacterium | 0 | 0 | 0 | 0 | 0 | 0 |
| Anaerostipes | 0 | 0.001929 | 0 | 0 | 0.002347 | 0.00179 |
| Anaerococcus | 0 | 0 | 0 | 0 | 0 | 0.003579 |
